# Whole-genome pre-amplification as a viable approach for genomic screening of FFPE-derived DNA samples

**DOI:** 10.64898/2026.03.26.714414

**Authors:** C. Guerrero Quiles, T. Lodhi, R. Sellers, S. Sahoo, J. Weightman, W. Breitwieser, D. Sanchez-Martinez, M. Bartak, A. Shamim, S. Lyons, K. J. Reeves, R. Reed, P. Hoskin, C.M. West, L. J. Forker, T.A.D. Smith, R. G. Bristow, D.C. Wedge, A. Choudhury, L.V. Biolatti

## Abstract

Whole-genome sequencing (WGS) enables comprehensive analysis of tumour genomes, but its use in formalin-fixed paraffin-embedded (FFPE) samples is limited by DNA fragmentation and low yields. Whole-genome amplification (WGA) methods such as multiple displacement amplification (MDA) can boost DNA availability but distort copy-number alteration (CNA) profiles. DNA ligation-mediated MDA (DLMDA) mitigates this bias by reconstituting fragmented templates, yet its performance in FFPE-derived DNA remains uncertain. We compared paired DLMDA pre-amplified (2h, 8h) and non-pre-amplified FFPE prostate tumour samples from 22 archival blocks (5, 15 and 20 years old). DLMDA increased DNA yield by 42- to 86-fold, with global CNA patterns largely preserved. However, DLMDA significantly reduced the number of detected CNA deletions and amplifications. These effects were independent of both block age and reaction time. CNA dropouts were randomly distributed across the genome, indicating that DLMDA does not introduce regional bias. Our results show that DLMDA enables robust DNA yield recovery and avoids false-positive CNA artefacts, but at the cost of reduced CNA sensitivity. While suitable for CNA screening pipelines through WGS, further improvements are required to minimise the false-negative risk and improve the technique’s sensitivity for FFPE-based genomics.

## Introduction

Whole-genome sequencing (WGS) is a powerful analytic technique for comprehensive studies of genetic alterations across the whole genome through untargeted sequencing^1^. In oncology, WGS has been used to characterise copy number alteration (CNA) changes and single-nucleotide variants (SNVs) to elucidate driver mutations and clinically informative molecular markers associated with patient prognosis and response to treatment^2,3^. Despite these advantages, routine clinical adoption is tempered by turnaround-time constraints, the uncertainty of the cost-effectiveness of undiscriminated WGS for each patient and the complexity of data storage and analyses. These limitations underpin the alternate use of targeted sequencing of validated markers as being more suitable for clinical implementation and patient stratification^1,4^.

Large reference cohorts have demonstrated the discovery power of WGS. The TCGA (The Cancer Genome Atlas)/ICGC (International Cancer Genome Consortium) Pan-Cancer Analysis of Whole Genomes analysed >2,700 samples across 38 tumour types, expanding the catalogue of non-coding drivers and structural events that shape cancer genomes. These analyses identified novel targets and biomarkers to improve cancer diagnosis and treatment^5,6^. The landmark consortia resources were assembled from prospectively collected fresh-frozen tissue^5,6^; yet, formalin-fixed paraffin-embedded (FFPE) samples dominate routine clinical pathology workflows^7^. Although FFPE samples preserve tumour DNA and are indispensable for retrospective translational studies, formalin fixation introduces DNA damage, fragmentation, cross-linking, and cytosine deamination, leading to poor yield, low-quality DNA samples. These can generate sequencing artefacts and reduce library complexity, increasing WGS error rates as compared to WGS using fresh-frozen material^7,8^.

Whole-genome amplification (WGA) is employed to increase DNA yield from limited and/or degraded inputs before high-throughput sequencing^9,10^. Several WGA techniques have been developed, with multiple displacement amplification (MDA) being the most widely used^9–11^. MDA uses the Φ29 DNA polymerase for isothermal strand-displacing synthesis, allowing high processivity and coverage of minute DNA quantities, successfully amplifying reduced DNA amounts with as little as 10 pg^9–11^. For genotyping applications, MDA-amplified DNA can achieve high SNV call concordance relative to unamplified DNA when analysing FFPE biopsies, supporting its utility for variant discovery when input is constrained^12^. By contrast, accurate copy-number profiling after MDA is more challenging: both empirical and comparative studies show that MDA introduces representation bias and allelic imbalance that can distort CNA landscapes and increase the false positive detection rate, particularly when starting from fragmented FFPE DNA^9,10,13^.

To mitigate MDA-associated bias in degraded templates, ligation-mediated repair strategies were proposed to reconstitute longer DNA fragments before amplification. In FFPE samples, ligation-based pre-treatments improved performance in comparative genomic array hybridisation experiments^14^, though their impact on next-generation sequencing (NGS)– based CNA calling remains unclear.

In many retrospective studies, the quantity of DNA extracted from FFPE blocks is frequently insufficient for WGS, particularly in older archival material or small biopsies, a problem frequently found in prostate cancer biopsies^15,16^. In such cases, pre-amplification is required to obtain enough DNA for WGS. Therefore, while unamplified DNA provides the most accurate CNA profiles when yields are adequate, DLMDA offers the potential to recover otherwise unusable samples to increase cohort sizes in retrospective studies. Here, we conduct a methodological comparison of paired DNA ligation-mediated MDA (DLMDA) pre-amplified versus non-pre-amplified FFPE-derived prostate tumour samples to evaluate the suitability of the technique for NGS-based CNA analysis.

## Methodology

### FFPE samples collection

Samples were retrospectively retrieved under ethics approval by the Manchester Cancer Research Centre Biobank Ethics Committee (Reference-18/NW/0092; Ref 293603860). Fifteen FFPE biopsies from patients with localised prostate cancer (T1–T2) were retrieved, comprising five samples aged 5 years, five aged 15 years, and five aged 20 years (Supplementary file 1). From each biopsy, a single 4 µm section was cut and stained with haematoxylin and eosin (H&E) for tumour content assessment by an expert pathologist. A minimum of ≥30% tumour tissue was confirmed in all samples before subsequent analyses.

### DNA extraction and DLMDA-based pre-amplification

Three 10 µm sections were obtained from each FFPE block. DNA was extracted using the QIAamp DNA FFPE Tissue Kit (Qiagen, Hilden, Germany; cat. no. 56404) according to the manufacturer’s instructions. DNA concentration was measured using the Qubit™ dsDNA Quantification Assay Kit (Thermo Fisher Scientific, Massachusetts, USA; cat. no. Q32851) following the manufacturer’s instructions.

DLMDA pre-amplification was performed using the REPLI-g FFPE Kit (Qiagen; cat. no. 150243). A minimum of 100 ng of total DNA in 10 µl DNase-free water was added to the FFPE ligation master mix, prepared according to the manufacturer’s protocol. Samples were incubated at 24 °C for 30 min to induce random ligation of fragmented DNA, followed by heat inactivation at 95 °C for 5 min. The REPLI-g Master Mix was then prepared according to the manufacturer’s protocol and added to the DNA-ligated samples. Samples were incubated at 30 °C for either 2 h or 8 h for pre-amplification. Reactions were terminated by incubation at 95 °C for 10 min.

### Whole-genome sequencing

100 ng of gDNA was sheared using 10 cycles of sonication on the Diagenode Bioruptor Pico (Diagenode, Liege, Belgium), followed by a further quantification and quality check. Indexed sequencing libraries were prepared from 50 ng of sheared DNA using the NEBNext Ultra II DNA Library Prep Kit for Illumina (New England Biolabs, Massachusetts, USA; cat. no. E7645L) with 4 cycles of amplification, according to the manufacturer’s instructions.

Libraries were sized and quality-checked using the Agilent Fragment Analyser (Agilent Technologies, Santa Clara, CA, USA) balanced by Qubit, and an equimolar pool was quantified by qPCR using the KAPA Library Quantification Kit (Roche, Basel, Switzerland; cat. no. 07960336001) for Illumina sequencing platforms. Paired-end sequencing was performed on an Illumina NovaSeq 6000 (Illumina, California, USA) with read lengths of 2×101bp.

### Data processing

Reads were quality checked and aligned to the GRCh38 reference genome using BWA (v0.7.7)^17^ with default options. Before downstream analysis, samtools (v1.2)^18^ was used to retain only uniquely mapped reads. BAM files were coordinate-sorted and de-duplicated via Picard (v1.96)^19^.

### Copy Number Analysis

HMMcopy (v0.99.0)^20^ was applied to the de-dupped BAM files to detect copy number change. The genome was divided into windows of fixed size (150 kb), and the read count was determined as the number of reads overlapping each window. GC content and mappability bias correction were performed on tumour and normal (if available) samples, filtering out GC content within the top or bottom 1 % quantile. The remaining windows with a mappability score greater than 0.9 were kept. The corrected read counts in each bin were used to calculate the log2ratio. A 6-state Hidden Markov model (HMM-based) approach was used to segment the data into regions of similar copy number profile and to predict a copy number alteration event (i.e., 0, 1, 2, 3, 4 or >5 copies of chromosome) for each segment. The fraction of the genome affected by copy-number changes (Genomic Instability Index, GII) was also calculated for each sample^20^.

### Statistical analysis

GII was calculated as the proportion of the genome with altered CN as defined by CNA. dGII (ΔGII) was calculated for 2h and 8h samples as the difference in GII between consecutive time points (2 h − 0 h and 8 h − 2 h, respectively). The Jaccard Index (JI) was calculated for 2h and 8h samples as the length of corresponding altered CN segments over the union of all altered CN segments between [0h vs 2h; 2h vs 8h]. Corrected JI values were calculated by subtracting predicted overlapping by random chance from the observed JI for each comparison. Statistical tests, where performed, are displayed and described within the figures.

## Results

### Pre-amplification increases DNA yields while largely preserving CNA profile of FFPE-derived DNA

To determine whether pre-amplification introduces systematic bias in CNA calling, we analysed 22 FFPE blocks: eight 5-year-old, nine 15-year-old and five 20-year-old. On average, 672.4 ng of DNA were retrieved per FFPE block (687 ng for 5-year-old, 852 ng for 15-year-old, 654 ng for 20-year-old) (Supplementary File 1). DLMDA was then performed for 2 h or 8 h on matched input samples to assess whether amplification time affects CNA representation. After 2h and 8h pre-amplification, DNA yield had an overall 41.9- and 85.6-fold increase, respectively (Figure 1).

**Figure 1.**
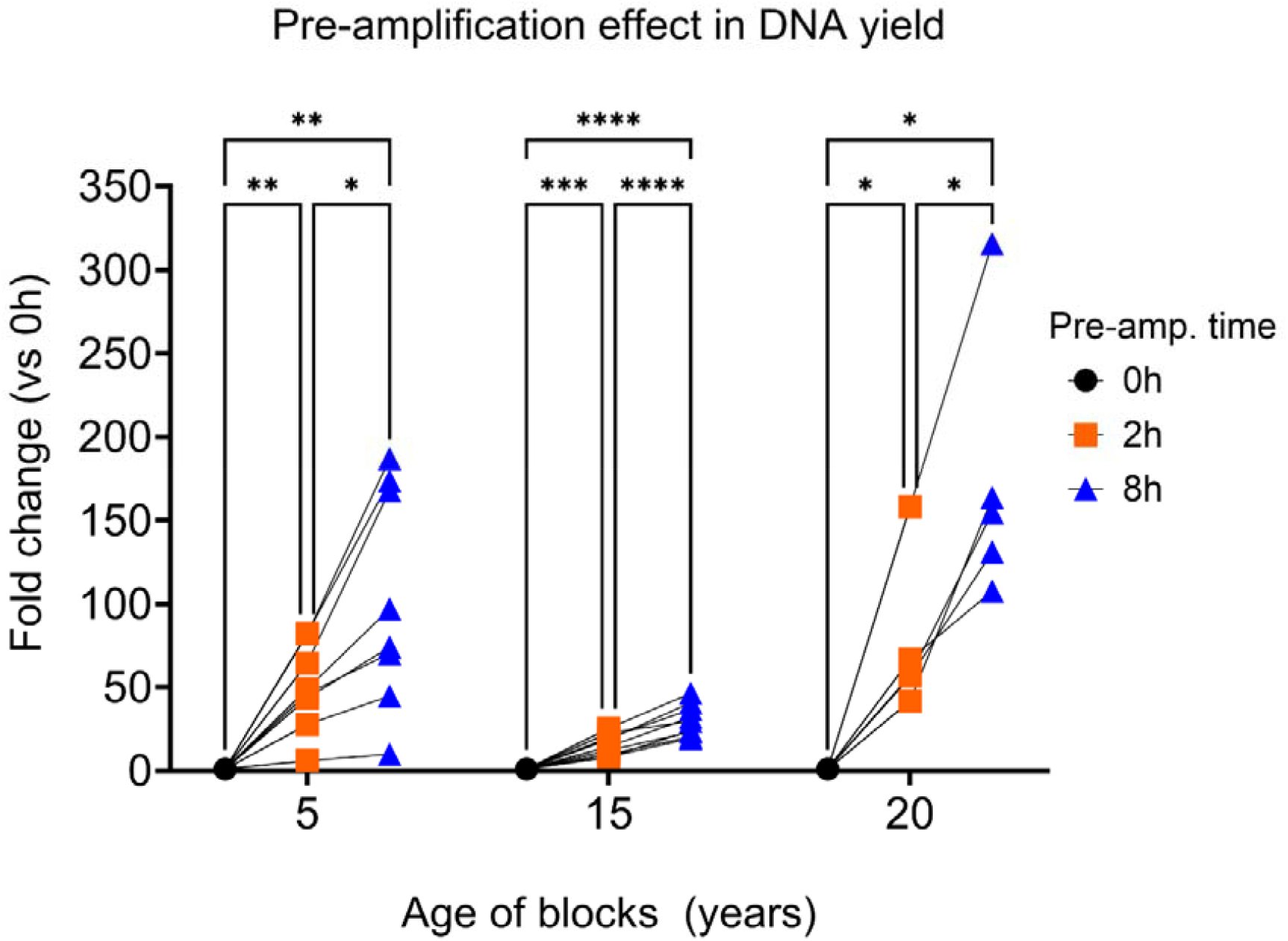
Pre-amplification increases the yield of DNA extracted from archival FFPE prostate tumour samples. The figure shows the relative fold-change increase in DNA yield after 2h (orange, squares) and 8h (blue, triangles) pre-amplification. Matched no pre-amplified samples (black, circles) are used as reference controls. Significance was determined using a two-way ANOVA with Tukey correction for paired samples analysis. Significance is represented as ^*^ for p adj.≤0.05, ^**^ for p adj. ≤0.01, ^***^ for p adj. ≤0.001, and ^****^ for p adj.<0.0001.

Across all conditions, global CNA patterns were conserved (Figure 2a–c), although CNA calling differences emerged between non-pre-amplified and pre-amplified inputs. Pre-amplified samples exhibited additional localised fluctuations, with increases (red) and decreases (blue) in CNA relative to unamplified DNA for both 2h (Figure 2b) and 8h (Figure 2c) pre-amplification. These fluctuations generally represented small-amplitude artefactual changes, although larger CNA disruptions after pre-amplification were observed in a small number of samples (Figure 2c). Overall, small distortions of the underlying CNA profiles were observed after 2h and 8h DLMDA.

**Figure 2.**
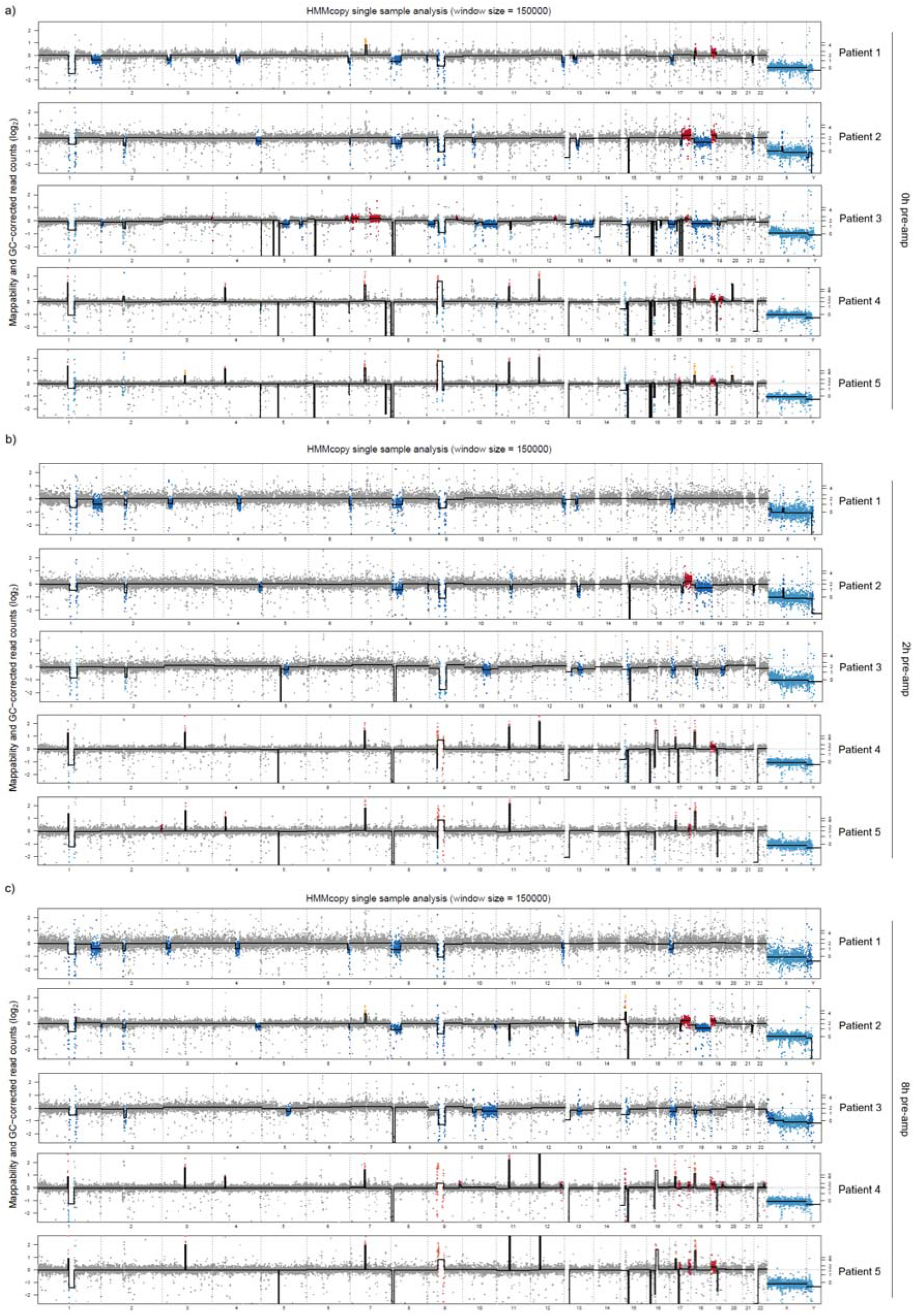
Pre-amplified prostate tumour FPPE samples have a distinct mapping of genomic regions. The figure shows increases (red) and decreases (blue) in CNA and their genomic location for a) non-pre-amplified DNA samples, b) 2h pre-amplified DNA samples and c) 8h pre-amplified DNA samples. The figure shows CNA changes due to pre-amplification of DNA samples derived from FFPE diagnostic biopsies. n=5 representative paired samples are shown across the experimental conditions.

### Pre-Amplification reduces CNA deletions and amplifications independently of FFPE block age

To dissect the contribution of specific CNA events to the observed reductions in GI, we independently quantified the frequency of CNA deletions and amplifications in pre-amplified versus non-pre-amplified samples.

Multivariate analyses showed a significant reduction in CNA deletions following pre-amplification, with decreases evident after 2 h (p≤0.01) and 8 h (p≤0.001), independently of FFPE block age (p=1) (Figure 3a). Grouped analysis of FFPE block ages corroborated this overall trend (p=0.0014), with statistical significance retained again only for 15-year-old blocks in stratified testing (Figure 3b).

**Figure 3.**
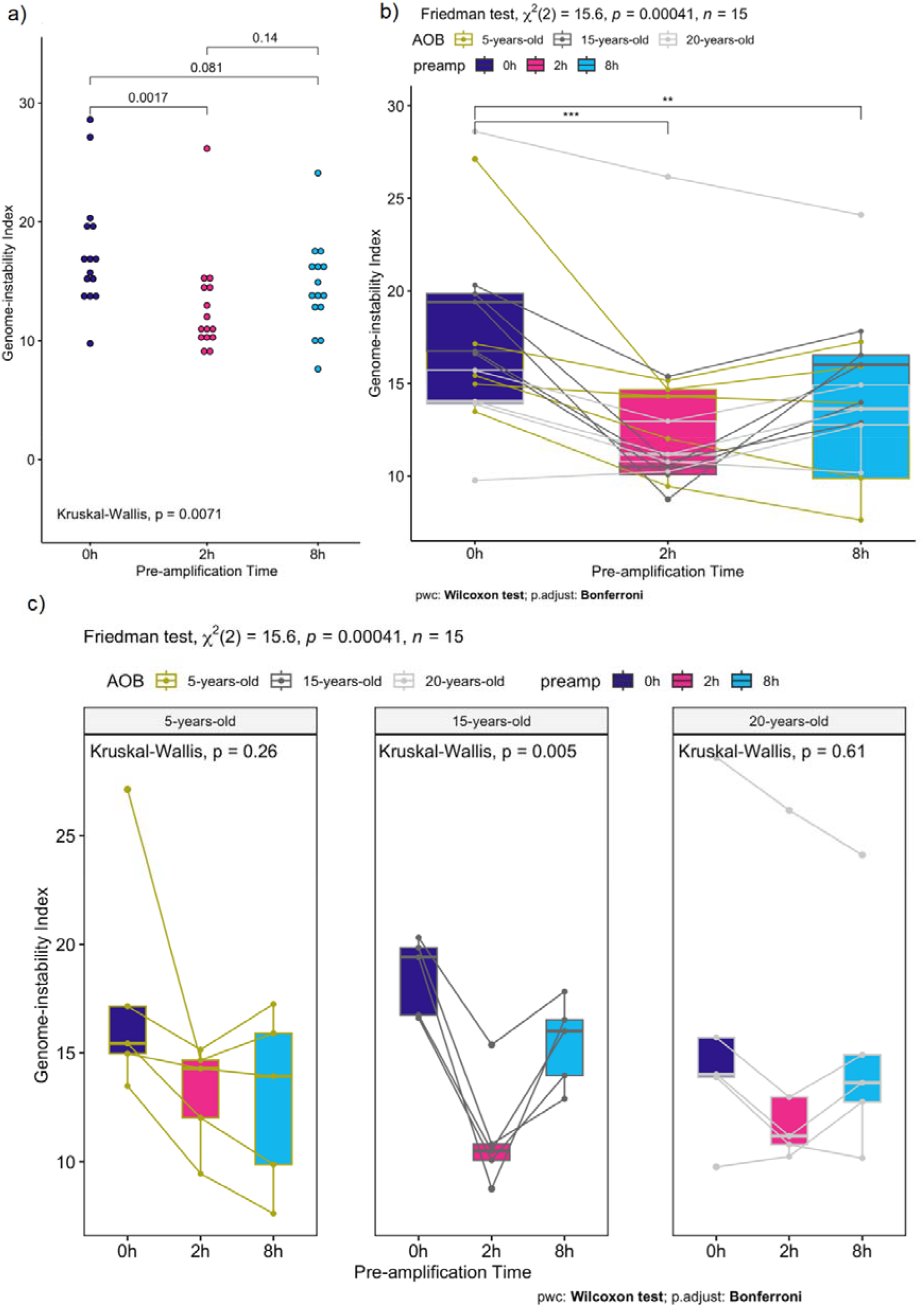
Pre-amplification reduces the genomic instability index (GII) independently of FFPE block age. (a) Unpaired analysis of whole-genome pre-amplified vs non-pre-amplified FFPE samples shows a GII decrease after 2h (p=0.0017) and 8h (p=0.081) pre-amplification. (b) Multivariate paired analysis confirmed significant GII decreases after 2h (p≤0.001) and 8h (p≤0.01) pre-amplification with non-confounding effect of the age-of-blocks (p=1). (c) Age-of-blocks grouped analysis confirmed a general overall decrease in GII after pre-amplification (p=0.00041). When analysed independently, significance was only found for 15-year-old blocks (p=0.005). GII is defined as the frequency of gene copy number variation (CNA) increases and decreases. Kruskal-Wallis evaluated overall significant differences for unpaired samples analysis and grouped paired samples analyses. Friedman test measures overall significant differences for paired samples analyses.

**Figure 4.**
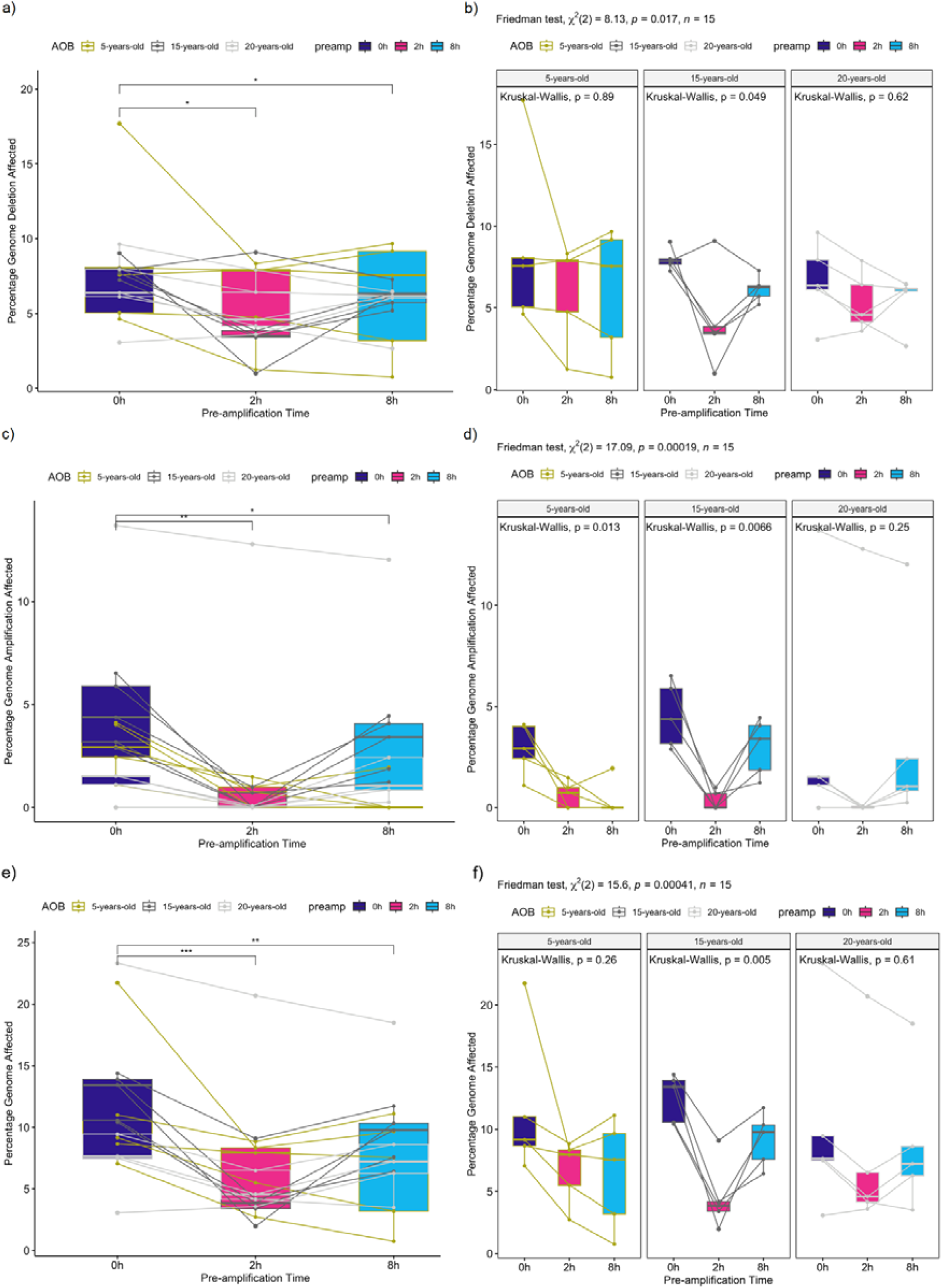
Pre-amplification decreases the number of identified genomic deletions and genomic amplifications. (a) Multivariate analysis shows pre-amplification decreases the percentage of identified gene copy number variation (CNA) deletions after 2h (p≤0.05) and 8h (p≤0.05), with no significant effect of the age-of-blocks (p=1). (b) Age-of-blocks grouped analysis corroborated an overall decrease in CNA deletions after pre-amplification (p=0.0017), with significance in independently grouped analysis found for 15-year-old blocks (p=0.049). (c) Multivariate analysis highlights a decrease in CNA amplification percentages after 2h (p≤0.01) and 8h (p≤0.05) pre-amplification, with no co-founding effect of age-of-blocks (p=1). (d) Age-of-blocks grouped analysis shows an overall decrease in CNA amplification percentages (p=0.00019), with grouped significant found for 5-year-old (p=0.013) and 15-year-old (p=0.0066) blocks. (e) A general decrease was observed in the percentage of identified deletions plus amplifications across the whole genome after 2h (p≤0.001) and 8h (p≤0.01) pre-amplification. (f) Age-of-blocks grouped analysis confirmed the overall combined decreased of identified deletions plus amplifications percentages (p=0.00041), with grouped significant found only for 15-years-old blocks (p=0.005). Friedman test measures overall significant differences for paired samples analyses. Kruskal-Wallis evaluated significant differences for grouped paired samples analyses.

Similarly, CNA amplifications were significantly reduced after pre-amplification, with decreases observed at both 2 h (p≤0.0001) and 8 h (p≤0.05) (Figure 3c). These reductions were again independent of block ages (p=1). FFPE blocks age-grouped analysis confirming overall significance (p≤0.0001), with the strongest effects detected in 5-year-old (p=0.013) and 15-year-old blocks (p=0.00073) (Figure 3d).

These results show that the decrease in GII observed after DLMDA is driven by a reduction in both CNA deletions and amplifications, regardless of FFPE block age.

### Pre-Amplification Does Not Introduce Regional Bias in CNA Detection

We investigated whether the CNA events undetected after pre-amplification were associated with specific genomic regions by calculating the JI, defined as the proportion of overlapping CNAs between pre-amplified and baseline samples.

Overall analysis revealed no significant differences in JI after 2 h and 8 h pre-amplification (Figure 5a). Age-of-blocks comparative analysis showed a proportional increase in JI for 15-year-old samples compared to 5-year-old samples (p=0.03), indicating higher concordance in older archival material (Figure 5b). Age-of-blocks grouped analysis showed no significant JI changes between 2h and 8h amplification within either the 5-, 15-, or 20-year-old FFPE samples (Figure 5c). The same analysis highlighted a reduced JI baseline following 2 h pre-amplification for 15-year-old FFPE biopsies compared to 5- and 20-year-old FFPE samples, providing context for the significant increase in overlapping CNAs observed for 15-year-old FFPE blocks following 8 h pre-amplification.

**Figure 5.**
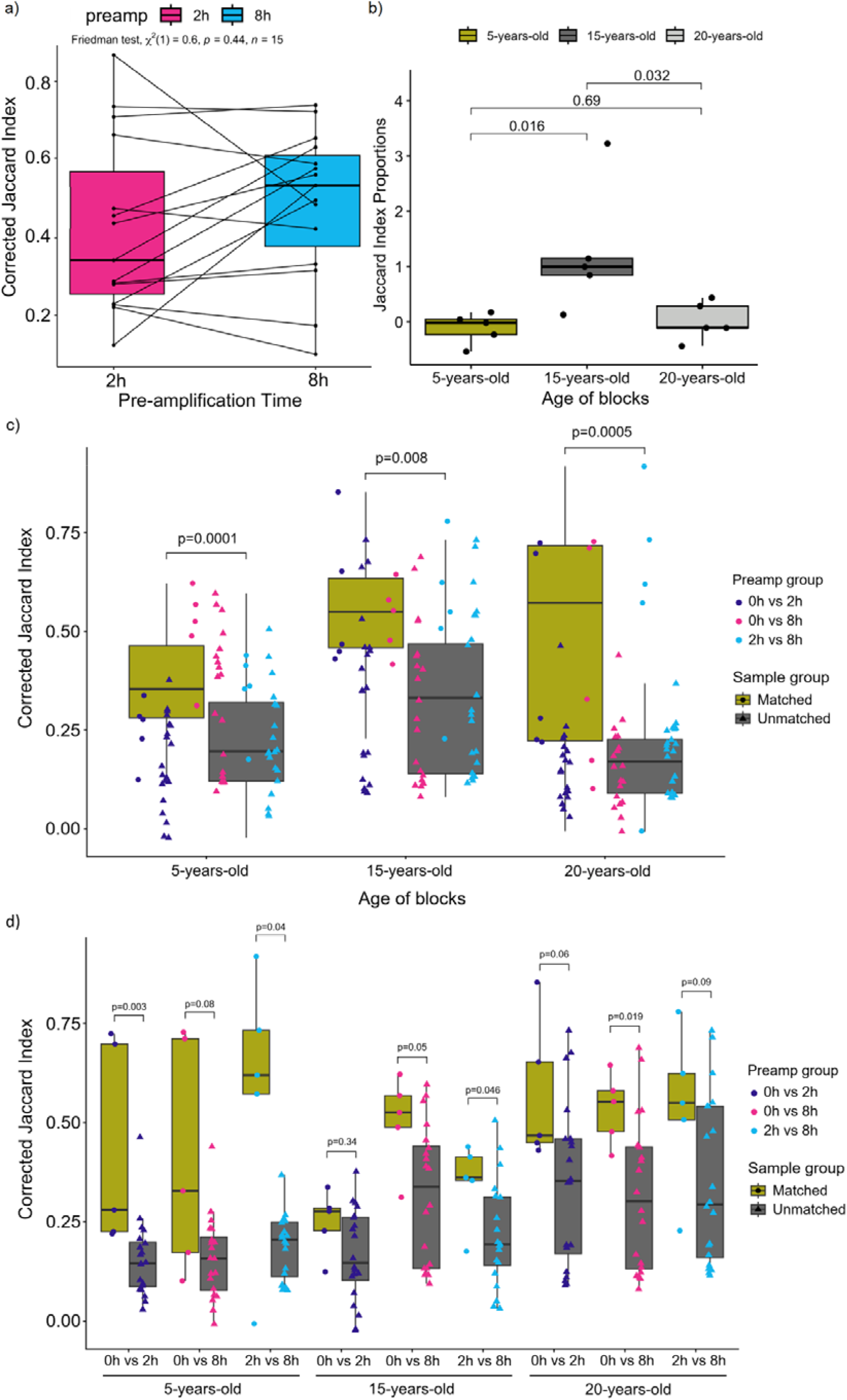
Genomic alterations undetected during pre-amplification are not associated with specific genomic regions. (a) Overall analysis shows no significant differences in Jaccard Index (JI) for pre-amplified samples. (b) Age-of-blocks analysis shows a proportional increase in JI for 15-year-old sample compared to 5-years-old samples (p=0.03). JI proportion was defined as the relative change in JI between 2h and 8h time points. (c) Multivariate analysis shows increased JI values for matched pre-amplified samples vs non-matched samples independently of age of blocks. (d) Age of blocks grouped analysis confirms higher JI values for matched vs unmatched samples, with significantly higher JI (p≤0.05) found for 0h vs 2h pre-amp (5-years-old), 0h vs 8h pre-amp (15-years-old, 20-years-old), and 2h vs 8h pre-amp (5-years-old, 20-years-old). JI is defined as the commonly occurring gene copy number variations (CNA) divided by all identified CNA within a group when compared to baseline (0h pre-amplification). JI proportion was defined as the relative change in JI between 2h and 8h time points. Friedman test measures overall significant differences for paired samples analyses. Kruskal-Wallis evaluated significant differences for grouped paired samples analyses.

For further evaluation, corrected JI values were calculated for matched (same patient) and unmatched (random matching, baseline) samples. Multivariate analysis showed consistently increased JI values for 5-(p=0.0001), 15-(p=0.008), and 20-years-old (p=0.0005) matched samples (Figure 5c). When independently analysed by age-of-blocks groups, higher JI was observed for matched samples across all conditions, being significant for 0 h vs 2 h pre-amp (5-years-old), 0 h vs 8 h pre-amp (15-years-old, 20-years-old), and 2 h vs 8 h pre-amp (5-years-old, 20-years-old) (Figure 5d). These results suggest that although pre-amplification reduces the absolute number of CNA deletions and amplifications, identified CNA alterations do not converge on specific genomic regions, supporting a largely random genomic distribution of these alterations following pre-amplification.

## Discussion

Here, we evaluated whether DLMDA can be applied to FFPE-derived tumour DNA to increase yield without compromising CNA profiling in WGS. Our results confirmed that DLMDA greatly increases DNA yield, achieving a 41–86-fold gain depending on incubation time. This fact is consistent with the high processivity of Φ29 polymerase^21^ and prior reports showing that MDA can be used to amplify limited DNA amounts^9,10,22,23^.

Our WGS comparison of pre-amplified and non-pre-amplified samples showed that DLMDA reduces both CNA deletions and amplifications, independently of FFPE block age. WGA has been previously linked to artificial CNA alteration and representation bias^9,10^. Xia et al. demonstrated that early WGA methods over-amplified genomic regions, generating artificial CNA amplifications and increasing the risk of detecting false positive targets^24^. Similarly, Pugh et al. observed both CNA increases and decreases after MDA in fresh tumour samples^13^. These studies suggested that WGA cannot be reliably used for CNA analysis without introducing false positive artefacts.

Our data refine this view by showing that DLMDA decreases both the number of detected CNAs deletions and amplifications, suggesting that DLMDA does not increase the false positive risk. In support of this concept, other MDA protocol improvements by Hammon et al. and Sidore et al. also reported increased DNA coverage and uniform amplification patterns, which closely resemble those of unamplified samples^22,23^. However, these previous studies evaluated the technique in fresh DNA derived from bacterial cultures (e.g. *E. coli*). Our study adds to the literature by evaluating the technique in archival FFPE prostate samples representative of the clinical setting, which have low DNA retrieval yields, and high DNA fragmentation levels^7,8^.

The capacity of MDA approaches to be used for CNA genomic screenings in FFPE-derived DNA has been challenged by the highly degraded level of the material^7,8^. MDA-associated errors in CNA quantifications are associated with the uneven amplification of the starting material, leading to the identification of false-positive targets^9,10,22^. Our observed reduction in GII after DLMDA support a previous study showing that DLMDA reduces sequencing artefacts caused by MDA amplification in FFPE-derived DNA^14^. Our study adds to the literature by showing DLMDA decreases GII and is suitable for CNA quantification through WGS screening pipelines.

A downside is the observed decrease in identified CNA, an effect also reported by Pugh et al^13^. It is possible that a small number of genomic sequences are not successfully ligated, impairing their later amplification and detection through WGS. This hypothesis warrants further investigation, as improvements in the DNA ligation step would reduce the detection of false negatives in CNS quantification through a DLMDA-WGS pipeline, improving the reliability of the approach. However, comparative analyses using equally pre-amplified samples as a reference baseline can reduce this risk, improving the approach’s reliability^13^. Despite DLMDA limitations, the findings suggest it can be used for CNA quantification in WGS screenings, with no increased risk of reporting false-positive targets.

In single-cell approaches, Bourcy et al. showed that longer MDA reactions are associated with enhanced pre-amplification bias^25^. However, we found no evidence of that risk when applying DLMDA to FFPE-derived DNA. This fact is relevant, as 8 h DLMDA greatly increased the DNA yield compared to 2 h and could be used for FFPE-derived samples of very low DNA yield.

Finally, a previously unexplored question was whether WGA-induced biases in CNA quantification can occur within specific genomic regions. We showed that CNA dropouts after pre-amplification occur randomly, rather than targeting specific regions. This observation implies that DLMDA does not distort the genomic map by disproportionately affecting informative loci. A related concern highlighted by Lasken et al. was MDA’s capacity to generate chimeric amplicons that could target specific genomic regions^26^. Further studies highlighted that those chimeric amplicons have no distribution preferences and do not generate data bias in subsequent analyses^10^.

Our study has limitations. We did not evaluate DLMDA’s capacity to induce bias in SNVs and allelic variant callings, although there is a strong consensus on MDA robustly preserving variant callings and MDA with a >98% precision^21–24,27–29^. This fact is based on the Φ29 DNA polymerase, used for MDA approaches, having high fidelity and proofreading capacity, which allows for preserving sequence integrity during WGA^21,27,28^. Although Saadi et al. reported small genotypic alterations in variant calling, they concluded that MDA methods are reliable, with results significantly correlating (r>0.99, p≤0.01) with those of unamplified samples^30^.

Another limitation is the lack of comparison with other WGA methods. Although MDA is more extended, Li et al. showed that MALBAC pre-amplification more reliably estimates CNAs alterations when compared to MDA in fresh tissue samples^31^. On the other hand, MDA was the most accurate approach when analysing allelic variants in the same study^31^. Furthermore, Del Rey et al. showed that MDA-based approaches are superior for WGS pipelines such as the one used in our study, accomplishing higher coverage and preserving allelic variants again using fresh tissue samples^32^. Despite the evidence, these facts have not been comparatively tested in FFPE-derived material and should be explored in future studies.

## Conclusion

Our results indicate that LMDA boosts DNA yields from archival FFPE samples in WGS workflows. We observe no significant increase in false positives CNA calls or evidence of region-specific amplification bias. DLMDA reduces the number of identified CNAs, which could hinder the discovery of relevant targets. This effect was independent of reaction length, suggesting longer reaction times can be used if required without additional downsides to the technique. Future work should improve the DNA-ligation efficiency to enhance CNA detection sensitivity. Additionally, comparison with alternative amplification methods is also required to place these findings in a broader technical context. Overall, our data suggests that DLMDA is suitable for CNA screening pipelines, with the main limitation of reduced CNA detection sensitivity. Further optimisation is still required to minimise the false-negative risk.

## Acknowledgements

We thank the Cancer Research UK Manchester Institute Histology Facility service staff for their support. Faculty of Biology, Medicine and Health, University of Manchester Biological Mass Spectrometry Facility (Bio-MS) staff for their support. Work was carried out at The University of Manchester, Cancer Research UK – Manchester Institute, and the Manchester Cancer Research Centre, which provided infrastructure support and access to core facilities for the experiments performed in this study.

## Author Contributions

Conceptualisation: C.G.Q., L.V.B., T.A.D.S, P.H., L.J.F., R.B., D.W., A.C., L.V.B; Methodology: C.G.Q., T.L., L.V.B, W.B.; Software: R.S., S.S.; Validation: C.G.Q., T.L.; Formal Analysis: C.G.Q., T.L., R.S., S.S., J.W., S.L., A.S.; Investigation: C.G.Q., T.L., R.S., S.S. W.B., S.L., J.W., M.B., D.S.M., L.V.B.; Resources: T.L., L.J.F., C.M.W., P.H., A.C.; Data curation: C.G.Q., T.L., K.R., R.R.; Supervision: C.M.W., P.H., R.B., D.W., A.C., L.V.B.; Funding acquisition: K.R., R.R., L.F., R.B., D.W., A.C.; Writing (original draft): C.G.Q.; Writing (review & editing): All authors

